# A scalable device-less biomaterial approach for subcutaneous islet transplantation

**DOI:** 10.1101/2020.06.26.172148

**Authors:** Alexander E. Vlahos, Ilana Talior-Volodarsky, Sean M. Kinney, Michael V. Sefton

**Author notes:** Corresponding author. (M.V.S.).

## Abstract

The subcutaneous space has been shown to be a suitable site for islet transplantation, however an abundance of islets is required to achieve normoglycemia, often requiring multiple donors. The loss of islets is due to the hypoxic conditions islets experience during revascularization, resulting in apoptosis. Therefore, to reduce the therapeutic dosage required to achieve normoglycemia, pre-vascularization of the subcutaneous space has been pursued. In this study, we highlight a biomaterial-based approach using a methacrylic acid copolymer coating to generate a robust pre-vascularized subcutaneous cavity for islet transplantation. We also devised a simple, but not-trivial, procedure for filling the cavity with an islet suspension in collagen. We show that the pre-vascularized site can support a marginal mass of islets to rapidly return streptozotocin-induced diabetic SCID/bg mice to normoglycemia. Furthermore, immunocompetent Sprague Daley rats remained normoglycemia for up to 70 days until they experienced graft destabilization as they outgrew their implants. This work highlights methacrylic acid-based biomaterials as a suitable pre-vascularization strategy for the subcutaneous space that is scalable and doesn’t require exogenous cells or growth factors.

**Summary:** In this study methacrylic acid copolymer coated tubes generated a robust vascular response in the subcutaneous space, which was critical to support islet transplantation in a streptozotocin-induced diabetic mouse model. More importantly, the subcutaneous pre-vascularization approach using this copolymer coating was scalable into a larger allogeneic rat model and returned animals to normoglycemia for up to 70 days. This platform highlights the potential of a scalable biomaterial approach for pre-vascularization of the subcutaneous space in larger animal models.

## 1.0 Introduction

The subcutaneous space is currently being pursued as an alternative transplant site for pancreatic islets, because it can support a large transplant volume (assuming a large area is feasible), is minimally invasive, and is accessible for graft retrieval in the event of complications^1,2^. However, the subcutaneous site is poorly vascularized and must be modified to support the engraftment of pancreatic islets. There have been various successful approaches for subcutaneously delivering pancreatic islets, such as delivering them within biomaterials^3^, with cells^1,4^, or in large macro devices^5,6^. These approaches all support islet engraftment; however, many require an excess of islets to achieve normoglycemia.

Alternative strategies to subcutaneously deliver pancreatic islets have been pursued, using a two-step approach to first prepare a cavity within the subcutaneous space, followed by islet transplantation into the preformed cavity^2,5,7^. These approaches focus primarily on reducing the initial inflammatory response, normally experienced upon transplantation and exploit the host’s foreign body response to materials. Even without additional measures these 2-step protocols have been shown to support the transplantation of a marginal mass of pancreatic islets^2,5,8^, making better use of a precious resource.

In some of these cases, the vascular nature of the foreign body response is specifically referenced in describing the tissue generated in the first step. This pre-vascularization enhances oxygen and glucose transport to the transplanted islets and enables them to rapidly respond to hyperglycemia, similar to the response when transplanted into the kidney capsule^5^. Different approaches for enhancing this pre-vascularization effect have been tested, such as delivering exogenous growth factors using biomaterials^9^. In comparison, others have embedded pancreatic islets within a microvascular mesh generated using fibrin and endothelial cells *in vitro*, before being subcutaneously transplantated^6^.

Therapeutic polymers such as methacrylic acid (MAA)-based biomaterials generate vessels when subcutaneously transplanted in a variety of different mouse models (CD1^10,11^, SCID/bg^3^ and C57BL6^12^), without the use of exogenous growth factors or cells. In a separate study, we showed that in coating a polypropylene mesh chamber with this copolymer^12^, we could restore diabetic C56Bl/6 mice to normoglycemia with syngeneic islets^13^; uncoated mesh chambers were ineffective in this model.

In this study, we adapted a method used by Pepper et al^2^. A silicone catheter was coated with a copolymer of MAA and isodecyl acrylate (IDA) and used to pre-vascularized the subcutaneous space in SCID/bg mice and Sprague Dawley (SD) rats. Following pre-vascularization, the catheter was removed, and pancreatic islets were delivered into the pre-vascularized cavity (**Fig. 1**). First, we determined when the MAA-coated tubes generated the maximal number of vessels after subcutaneously implanted in immune-compromised SCID/bg mice. We confirmed that the pre-vascularized site could support the engraftment of a marginal dosage of pancreatic islets. Finally, we showed that this MAA-based approach could be scaled up into a larger rat model and support the long-term engraftment of allogeneic islets in immunosuppressed recipients. This study highlights the potential of using a biomaterial-based approach to pre-vascularize the subcutaneous space for islet transplantation, with emphasis on its ability to be scaled up into larger animal models.

**Figure 1:**
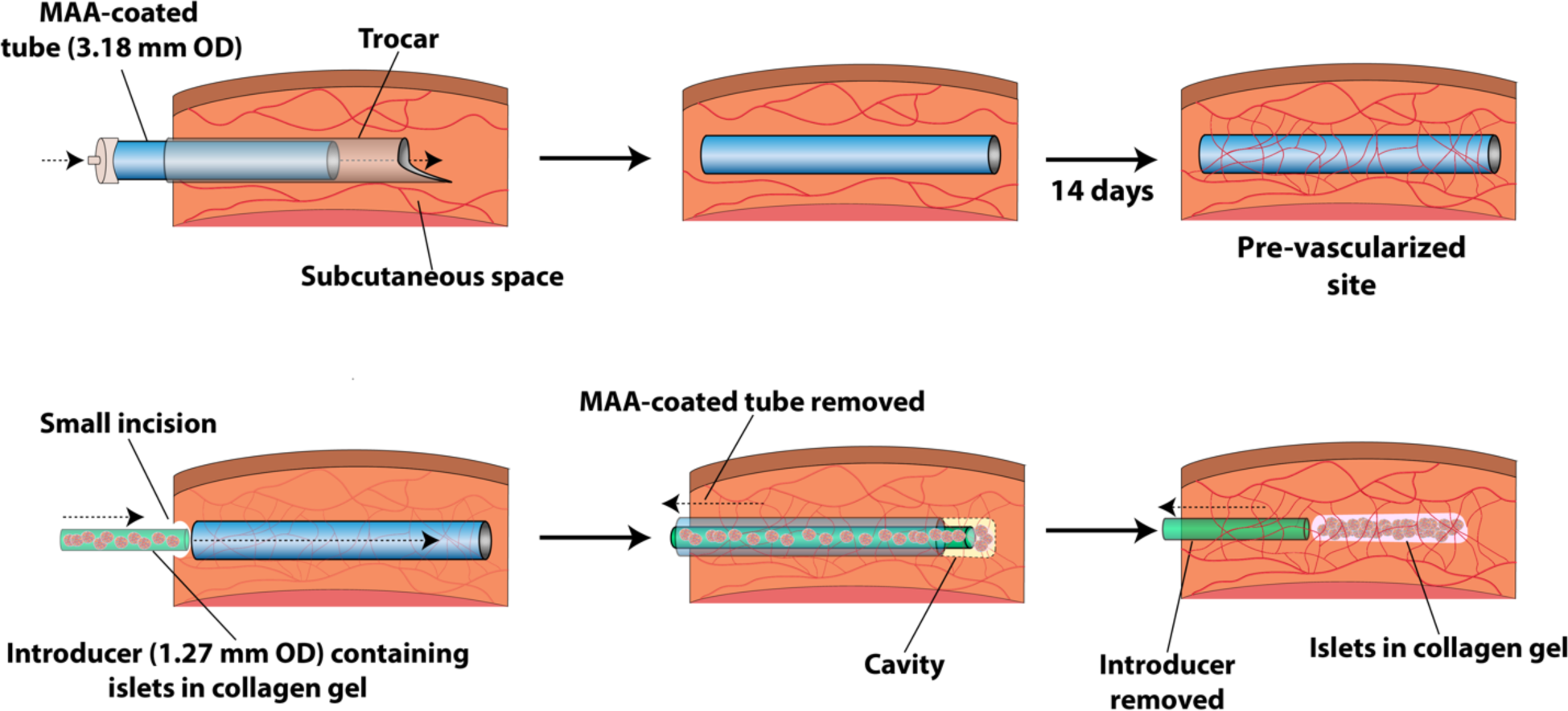
MAA-coated tube for pre-vascularization of the subcutaneous space and islet transplantation in mice or rats. MAA-coated silicone tubes (or uncoated tubes) were inserted into the subcutaneous space using a trocar. After 14 days when the space around the MAA coated tube became vascularized, a small incision exposed the MAA-coated tube and an introducer catheter containing the islets in the collagen gel was inserted through the MAA-coated tube into the pre-vascularized space. As the introducer was inserted, the MAA-coated tube was slowly removed to allow the islets in the collagen gel to be deposited into the pre-vascularized cavity. After the islets in the collagen gel were delivered, the introducer was removed, and the incision was sutured. The tubing length was 3 cm and 6 cm in mice and rats, respectively. The available transplant volume was dependent on the length of tubing as summarized in **Supplementary Table 1**.

## 2.0 Materials and Methods

### 2.1 MAA co-polymer coating of silicone tubes

Methacrylic acid (MAA, Sigma) was co-polymerized with isodecyl acrylate (IDA, Sigma) as previously described^11,14^. Silicone tubes (1.57 mm inner diameter (ID) x 3.18 mm outer diameter (OD), BSIL-T062, Instech) were dip coated twice with purified 40% mol MAA co-polymers (dissolved in THF (50 mg/mL)) and allowed to air-dry overnight under sterile conditions. Once the tubes were completely dry, they were gas sterilized with ethylene oxide prior to transplantation. Uncoated silicone tubes were also gas sterilized with ethylene oxide prior to transplantation and used as controls.

### Transplantation of MAA-coated tubes for pre-vascularization

All animal experiments and surgeries were performed at the University of Toronto and were approved by the Faculty of Medicine Animal Care Committee. 5-6 week SCID/Bg mice (Charles River) or 175-200 gram Sprague Daley Rats (Charles River) were subcutaneously implanted with MAA-coated silicone tubes (3 cm or 6 cm long, respectively). Uncoated silicone tubes were used as controls. Briefly, following anaesthetization and administering ketoprofen (0.2 mg/kg), a small incision was made on the skin close to the upper dorsum. Using a hematostatic clamp, blunt dissection was performed to create a tunnel, into which a MAA-coated tube or uncoated tube control was inserted. The incision was sutured using Polysorb™ 3-0 sutures (Covidien) and cleaned using betadine. Non-diabetic animals were sacrificed on days 14 and 21 and implants were harvested for histological analysis. Alternatively, tubes in diabetic animals were replaced with islets in collagen gel following a protocol described in the next section.

### 2.2 Islet Isolation

Primary rat islets were isolated from 12-week Wistar Rats (Charles River) as previously described^1,4^. Briefly, the pancreas was perfused using a collagenase solution (Cizyme, Vitacyte), which was then excised and incubated in a shaking water batch at 37°C for 19 minutes. Following mechanical digestion and density purification using Histopaque 1100 (Sigma Aldrich), pancreatic islets were handpicked and islet equivalent units (IEQ) were calculated based on volumetric assumptions. Pancreatic islets were cultured in RPMI-1640 (Gibco) supplemented with 10% FBS (Gibco) and 1% PenStrep (Gibco). For transplantation studies, multiple isolations were performed to procure enough islets to span multiple test groups.

### 2.3 Islet transplantation

SCID/bg mice were made diabetic using a single i.p. injection of streptozotocin (180 mg/kg, Sigma Aldrich) one week before islet transplantation (**Supplementary Figure 1A**). Sprague Daley rats were made diabetic using a single i.v. injection of streptozotocin (40 mg/kg) via the tail vein one week before islet transplantation. Animals were monitored daily (OneTouch UltraStar2) and only animals with non-fasting blood glucose levels above 20 mM for two consecutive days were used for transplantation studies.

Prior to transplantation, the pancreatic islets were washed three times with PBS and then mixed with neutralized Type I collagen (3.0 mg/mL – PureCol, Advanced Biomatrix). Depending on the recipient species, different numbers of IEQs were mixed with various volumes of collagen as outlined in **Table 1**. For mouse implants, human umbilical vein endothelial cells (HUVEC, Lonza) were added to the islet/collagen mixture at a dosage of 10^6^ HUVEC/mL of collagen; HUVEC had been previously observed to improve retrievability of collagen-based islet transplants^1^. We used HUVEC here to enable a comparison with earlier studies. The islet/collagen mixture was drawn into PE90 tubing (0.86 mm ID x 1.27 mm OD, IntraMedic) and gelled at 37°C for 1 hour.

**Table 1:** Composition of islet/collagen grafts for transplantation into pre-vascularized subcutaneous space. Details relating to the tube length, and available transplant volume can be found in **Supplementary Table 1**.

| Recipient | Number of IEQs per recipient | Volume of neutralized collagen (μL) |
| --- | --- | --- |
| SCID/bg mice (~20-25 g) | 250 | 20 μL |
| Sprague Daley rats (~325 - 375 g) | 4000 | 110 μL |

To access the subcutaneously inserted MAA-coated or uncoated tubes, an incision was made close to the upper dorsum of the anaesthetized animals. The tube was flushed 3 times with PBS, and residual PBS was then aspirated using an 18-G syringe. The PE90 tubing containing the islet/collagen mixture was inserted through the subcutaneously placed tube, which was used as a guide to inject the islet/collagen mixture into the pre-vascularized site. The larger tube was partly removed during the injection (**Fig. 1**). Following the injection, both tubes were removed, and the incision was then sutured using PolysorbTM 3-0 sutures (Covidien). Age-matched mice were used as no pre-vascularization controls, delivering the islet/collagen mixture (with HUVEC) using the PE90 tubing into the unmodified subcutaneous space.

Xenogeneic islet transplantations into the SCID/bg mouse did not require any immunosuppression. For allogeneic rat islet transplantations (**Supplementary Figure 1B**), the recipients were immunosuppressed using a combination of anti-lymphtocyte serum (ALS, Lot: J2445 and J2546, Accurate Chemical Scientific Corporation), mycophenolic acid (Myfortic, Novartis) and FTY-720 (fingolimod, Biorbyt). ALS was administered as a single i.p. injection (0.5-0.75 mL, dosage determined by lot) 3 days prior to allogeneic islet transplantation. Starting on the day of islet transplantation and daily for the entire duration of the experiment, rats were administered MMF (20 mg/kg, which was tapered to 0 mg/kg between day 14 to 21) and FTY-720 (2 mg/kg) via oral gavage. For long term rat studies, an additional injection of ALS was given at day 21 post transplantation.

For mice, the non-fasting blood glucose (NFBG) levels were monitored until day 28 for animals in the unmodified subcutaneous space and day 42 for animals in the MAA-coated or uncoated groups. For rats, NFBG was monitored until day 21, or day 90 for long-term studies. At experimental endpoints implants were removed and processed for histology.

### 2.4 Glucose tolerance test

SCID/bg mice were fasted 4 hours prior to being administered an intraperitoneal glucose tolerance test (IPGTT) on days 7 and 14 post transplantation. Animals were given an intraperitoneal injection of glucose (2 mg/kg) and then blood glucose was monitored 15, 30, 60 and 120 minutes post injection via tail-vein blood sampling.

Sprague Daley rats were administered an oral glucose tolerance test (OGTT) on days 14, and 85 post-islet transplantation. Rats were fasted overnight and then administered a glucose solution (2 mg/kg) via oral gavage. Animals were monitored at 15, 30, 60 and 120 minutes post oral gavage by tail-vein blood sampling. An area under the curve analysis was performed to compare among the different groups.

### 2.5 Histological Analysis

Implants with and without pancreatic islets were harvested and fixed using 4% paraformaldehyde. Implants were processed at the Pathology Research Laboratory at the Toronto General Hospital. Samples were cut 100 μm apart and serially stained for Masson’s trichrome, hematoxylin and eosin (HE), CD31, smooth muscle actin (SMA), and insulin. Sections were scanned (20X, Aperio ScanScope XT; Leica Microsystems, Concord, ON, Canada) at the AOMF Imaging Facility (UHN, Toronto). Scans were analyzed using ImageScope software version 11 (Aperio). For islet transplantation studies, only sections that had insulin positive staining were used to quantify vessel counts.

### 2.6 Statistical Analysis

A one-way ANOVA with a Games-Howell post-hoc test was used to compare the means among groups, unless otherwise stated. Data were considered statistically significant at a p value of 0.05. All statistical analysis was performed using SPSS software version 22 (IBM).

### 2.7 Data availability

The datasets generated during and/or analyzed during the current study are available from the corresponding author on reasonable request.

## 3.0 Results

### 3.1 Pre-vascularization using MAA-coated silicone tubes

Firstly, we wanted to validate the angiogenic effect of the MAA coating by measuring vessel densities after subcutaneous transplantation into SCID/bg mice. At day 14, the subcutaneous space around the MAA-coated tubes had a significantly greater number of vessels than the uncoated tube control (**Fig. 2 –** 193 ± 22 vs. 76 ± 17 CD31^+^ vessels/mm^2^, p < 0.01). However, at day 21, this difference was no longer statistically significantly (**Fig. 2 –** 131 ± 27 vs. 87 ± 26 CD31^+^ vessels/mm^2^). Therefore, subsequent islet transplantation studies were performed 14 days after inserting the MAA-coated tubes to maximize the number of vessels present at the time of islet transplantation.

**Figure 2:**
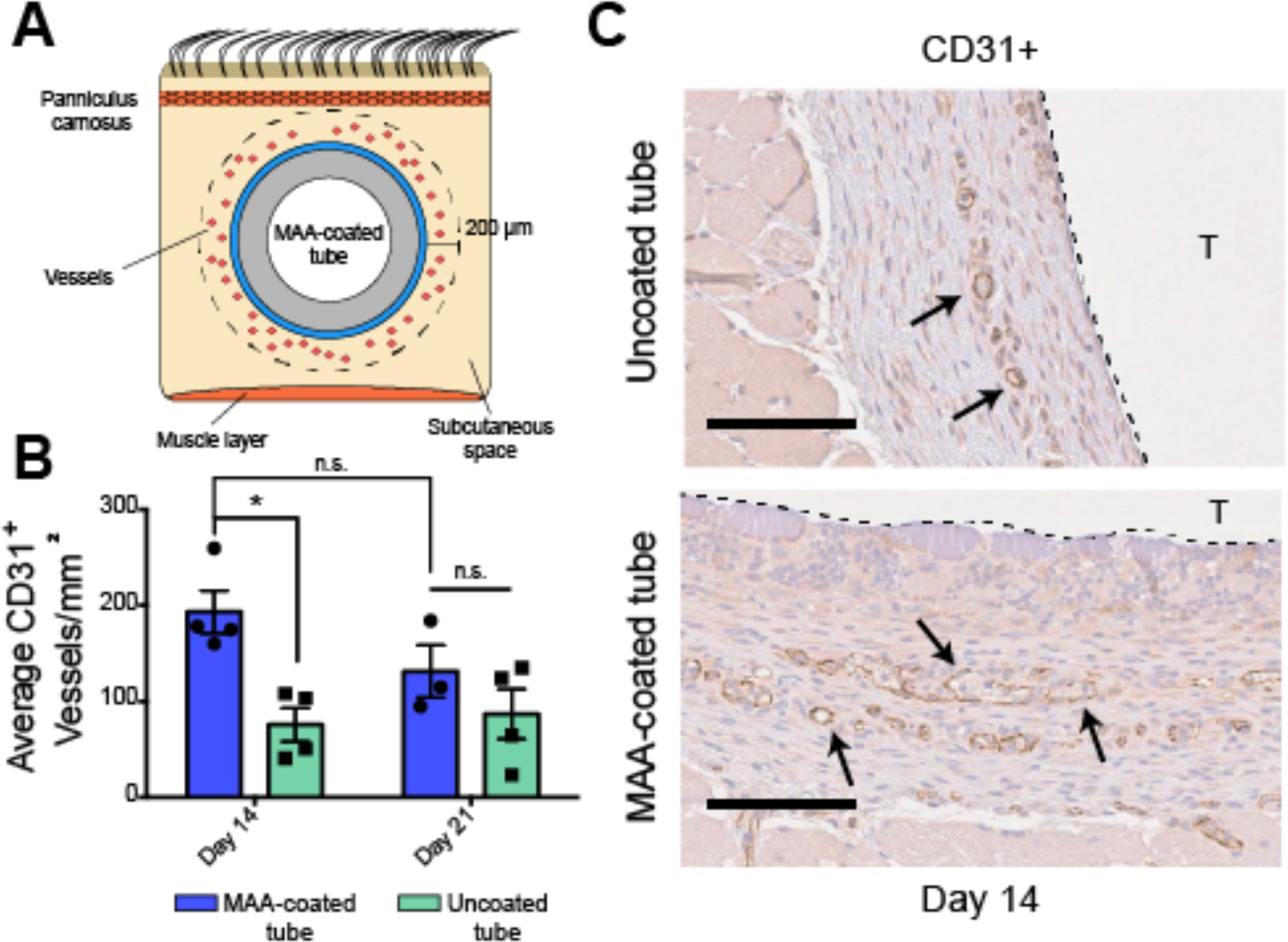
MAA-coated tubes vascularized the subcutaneous space of SCID/bg mice. **A)** Cross-sections of day 14 explants were used to calculate the average vessel densities in a 200 μm ring around the tube. **B)** The subcutaneous tissue of animals subcutaneously implanted with a MAA-coated tube (n = 3) had significantly more CD31^+^ vessels in the surrounding area compared to animals transplanted with an uncoated silicone tube (n = 4) at day 14 (one-way ANOVA, p < 0.05). At day 21, vessels began to regress in the MAA-coated tube group, albeit this difference was not statistically significant relative to day 14 counts. In addition, this difference in the number of vessels in the MAA-coated group was not statistically significant when compared to the uncoated controls at day 21. There was no significant difference in the number of vessels within the uncoated silicone group between days 14 and 21. **, p < 0.01. Average +/- SEM. **C)** Representative histological images of CD31^+^ vessels (arrows) in the subcutaneous space surrounding an uncoated silicone tube (top panel) or a MAA-coated tube (bottom panel). The dotted line denotes the boundary of the tube (T) with the subcutaneous space. Scale bars = 100 μm.

### 3.2 Subcutaneous delivery of pancreatic islets into the pre-vascularized space

The next step was to determine if the vessels created by the MAA-coated tube would support a marginal mass of pancreatic islets. Rat pancreatic islets (250 islet equivalent units - IEQ) within a collagen gel (mixed with 1 million HUVEC per mL of collagen) were injected into the subcutaneous cavity created by the MAA-coated tube or the uncoated silicone tube (**Fig. 3A – green and blue lines**). These transplanted islets were sufficient to return STZ-induced diabetic SCID/bg to normoglycemia (**Fig. 3B)** – in 3 of 5 MAA-coated tubes and in 4 of 5 uncoated silicone tubes. Age-matched SCID/bg mice with unmodified subcutaneous spaces (i.e., without pre-vascularization) were also used as recipients as a no pre-vascularization control group, and only one animal (of five) transplanted with pancreatic islets returned to normoglycemia (**Fig. 3A – red line**). The average time to return to normoglycemia was comparable between the MAA-coated and uncoated silicone tube groups.

**Figure 3:**
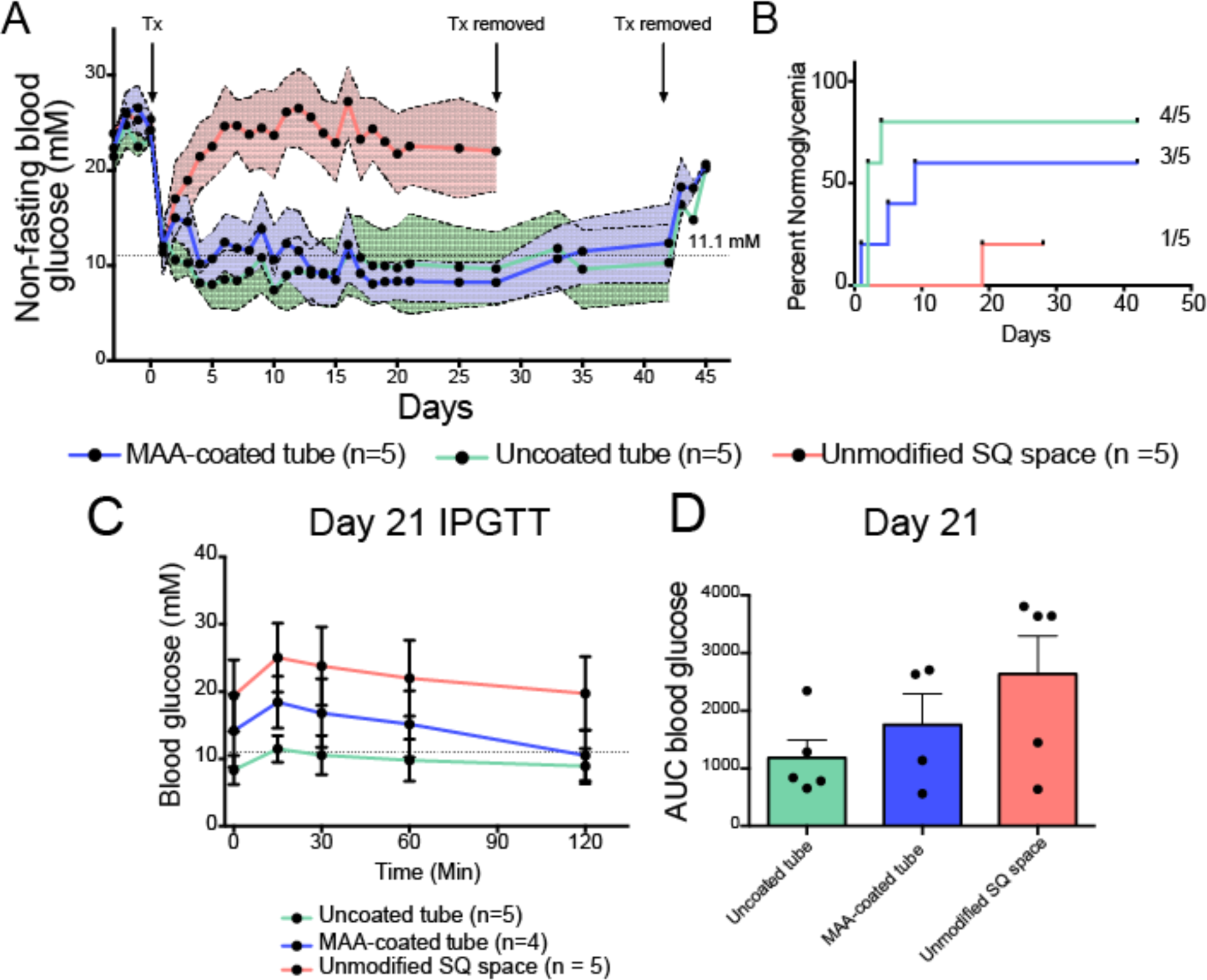
250 IEQ mixed with HUVEC and collagen implanted into the pre-vascularized subcutaneous space of SCID/bg mice. **A)** Average non-fasting animals of animals whose subcutaneous space was either pre-vascularized using a MAA-coated tube (blue line) or an uncoated silicone tube (green line) or left unmodified (red line). Pre-vascularized animals using a MAA-coated tube or an uncoated silicone tube rapidly returned to normoglycemia (< 11.1 mM) following islet transplantation. Animals that had become normoglycemic by day 42 had their implants removed and showed a return to hyperglycemia, however some explants (1 of 3 for MAA, and 3 of 4 for uncoated) could not be fully removed. Individual non-fasting blood glucose levels are in **Supplementary Figure 5**. The shaded areas around the lines represent the standard error of the mean for the respective groups. **B)** Kaplan-Meier curve indicating the day animals returned to normoglycemia and the proportion of animals that returned to normoglycemia in the respective groups. **C)** An IPGTT was performed at day 21 and only animals in the MAA-coated and uncoated groups returned to normoglycemia after being challenged with the glucose bolus. The animals that had islets transplanted into the unmodified subcutaneous space were intolerant to the glucose bolus as revealed by the **D)** AUC analysis. n = 5 for all groups. Average +/- SEM.

On day 21, animals were fasted for 4 hours before being administered a glucose bolus via intraperitoneal injection (IPGTT). Animals from both the MAA-coated and uncoated silicone tubes were able to respond to the glucose bolus and return to normoglycemia within 120 minutes (**Fig. 3C – blue and green lines**). Animals that had islets transplanted into the unmodified subcutaneous space were unable to respond to the glucose bolus and remained hyperglycemic (**Fig. 3C – red line**).

Since animals in the unmodified subcutaneous group were still hyperglycemic and could not respond when administered a glucose bolus, they were sacrificed, and their implants were removed at day 28. Animals in the MAA-coated and uncoated silicone tube groups were monitored for longer and sacrificed at day 42. Animals that had returned to normoglycemia had their grafts removed without sacrificing the animals and levels returned to hyperglycemia (**Fig. 3A**) – however not all of the grafts could be reliably retrieved. Pancreatic islets were structurally intact in both the MAA-coated and uncoated silicone tubes, but islets were fragmented in the unmodified SQ space group (**Supplementary Figure 2A**).

Retrieved grafts were well vascularized in all three groups (**Supplementary Figure 2A, left panels**), however the fragmented islets within the unmodified SQ space group did not have intra-islet vessels. The average vessel density was highest in the grafts retrieved from the MAA-coated tube group (105 ± 15 vessels/mm^2^), which was significantly higher than the unmodified SQ space group (54 ± 11 vessels/mm^2^). This value was also higher than the uncoated silicone tube group (60 ± 10 vessels/mm^2^) but this was not significant (**Supplementary Figure 2B**).

### 3.3 MAA-based vascularization of rats

The next step was to determine if the MAA-coated tubes would also generate a pre-vascularized cavity when deployed in the Sprague Daley (SD) rat model. Since we observed a significant difference in the number of vessels at day 14 in the SCID/bg model, we selected that time point for pre-vascularization in the SD rat model. Indeed, MAA-coated tubes had a significantly greater number of CD31^+^ vessels (**Fig. 4B –** 370 ± 29 CD31^+^ vessels/mm^2^) compared to the uncoated silicone tube control (204 ± 22 CD31^+^ vessels/mm^2^) at day 14 (**Fig. 4A**, p < 0.01). In addition, vessels generated in both groups were observed to be SMA^+^ (**Fig. 4C**), suggesting that these vessels were mature.

**Figure 4:**
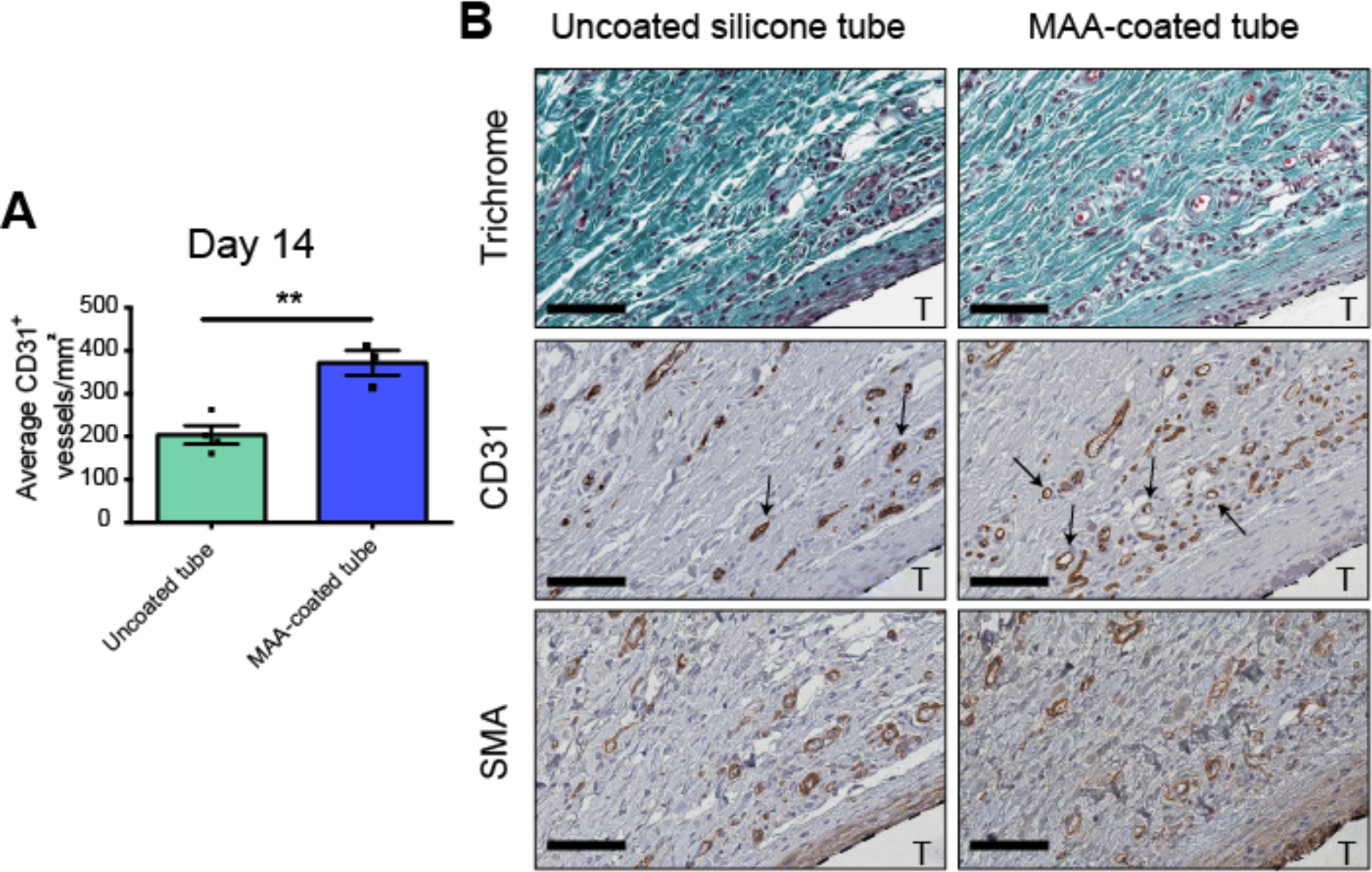
MAA-coated tubes pre-vascularized the subcutaneous space of Sprague Dawley Rats. **A)** The surrounding subcutaneous space of rats implanted with the MAA-coated tube had significantly more CD31^+^ vessels compared to the uncoated tube (p < 0.01). Average +/- SEM, n = 4 for both groups. **B)** Representative histology images of the surrounding subcutaneous space 14 days after an uncoated tube or a MAA-coated tube was inserted. Both conditions had mature vessels in the surrounding subcutaneous area (arrows - CD31^+^ and SMA^+^). T = tubes, Scale bars = 100 μm. ** p < 0.01.

### 3.4 Allogeneic islet transplantation into rats

After confirming the angiogenic benefit of the MAA-coating, outbred SD rats were subcutaneously transplanted with allogeneic Wistar rat islets. Unlike the immune-compromised SCID/bg mice, SD rats must be immunosuppressed to support allogeneic islet transplantation. For this study, SD rats were administered an immunosuppression regimen consisting of anti-lymphocyte serum (ALS), fingolimod (FTY-720), and mycophenolic acid (MMF) (**Supplementary Figure 1B**).

STZ-induced diabetic SD rats were transplanted with 4000 islet equivalent units into the pre-vascularized cavity created using MAA-coated tubes, uncoated silicone tubes, or into age-matched non pre-vascularized recipients. As expected, SD rats that were pre-vascularized with MAA-coated tubes quickly returned to normoglycemia (< 11.1 mM non-fasting blood glucose in 8 of 11 recipients) when transplanted with allogeneic islets (**Fig. 5A – blue line**). In comparison, only two of six rats in the group that were pre-vascularized with the uncoated silicone tube also returned to normoglycemia (**Fig. 5A – green line**), and none of the animals without pre-vascularization returned to normoglycemia (**Fig. 5A – red line**). Animals that were pre-vascularized showed a significantly faster return to normoglycemia compared to the unmodified subcutaneous group (p < 0.05, **Fig. 5C)**.

**Figure 5:**
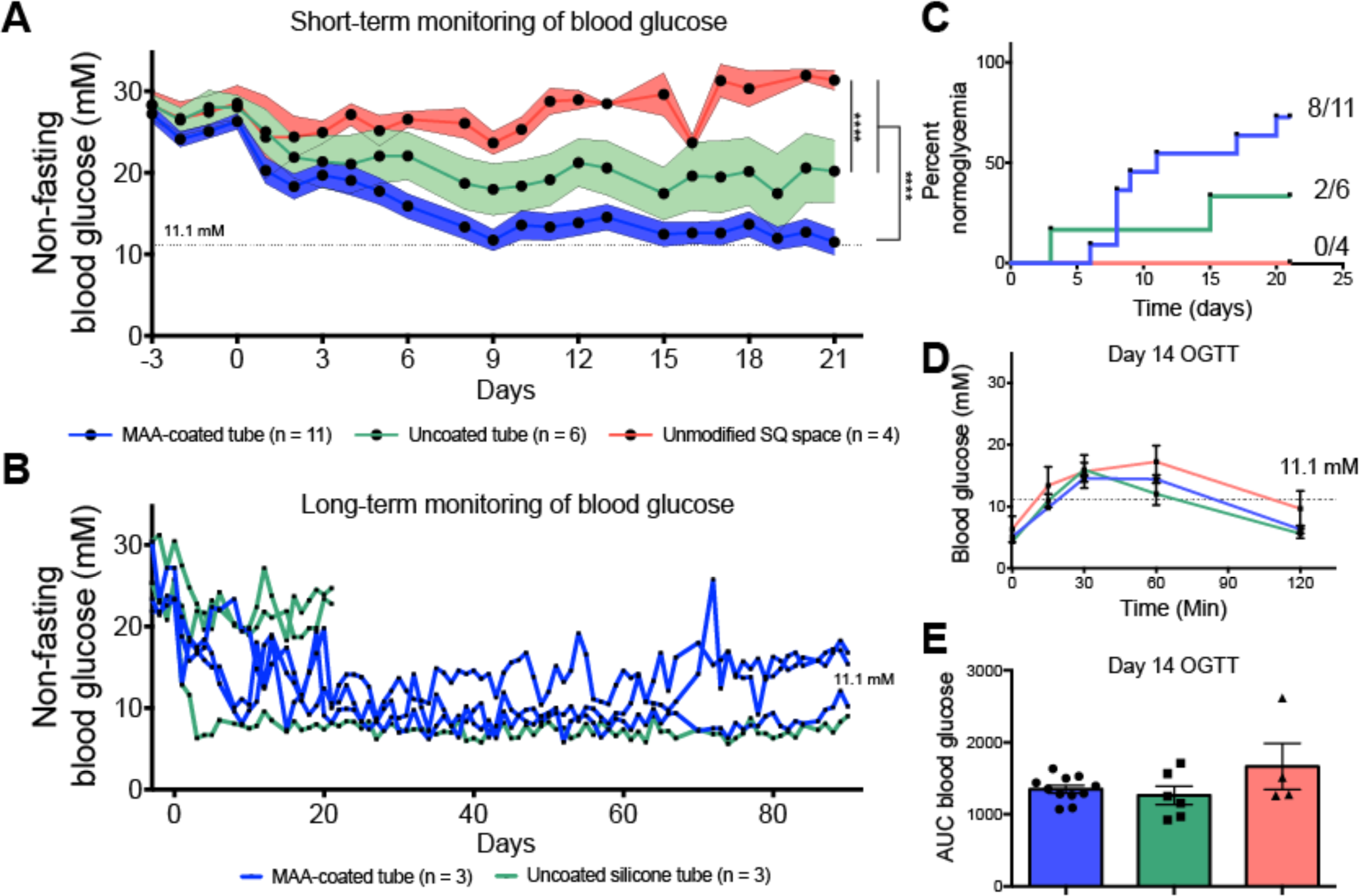
Allogeneic islets transplanted into pre-vascularized cavity created with MAA coated tube returned diabetic SD rats to normoglycemia. **A**) Average non-fasting blood glucose levels of all animals in the study that were transplanted with 4000 IEQ into a pre-vascularized space created with a MAA-coated tube (blue line) or an uncoated silicone tube (purple line), or into the unmodified subcutaneous space (no pre-vascularization, red line). Animals transplanted with the MAA-coated tube had significantly lower non-fasting blood glucose readings compared to the unmodified subcutaneous control and the uncoated silicone tube (p < 0.001, repeated-measures ANOVA). **B)** Long-term graft efficacy was also assessed in the MAA-coated and uncoated silicone tube groups, where animals remained normoglycemic until approximately day 70 when grafts begun to show destabilization. Two of three animals in the uncoated silicone groups were sacrificed at day 21 due to poor glycemic control attributed to graft failure. **C)** Kaplan-Meier curve showing when animals became normoglycemic and the relative number of animals that returned to normal. Rats in the MAA-coated tube group returned to normoglycemia significantly faster than animals in the no pre-vascularization group (* p = 0.0290, log-rank Mantel-Cox test). **D)** On day 14, all rats were fasted overnight and administered an oral glucose tolerance test. Animals from all groups were responsive to the glucose bolus and returned to normoglycemia within 2 hours. There were no significant differences observed among the groups, which was confirmed via **E)** AUC analysis. n = 11 for MAA-coated group, n = 6 for uncoated silicone group and n = 4 for the no pre-vascularization control group.

The majority of rats in this study had their grafts assessed for up to 21 days, but some animals in the MAA-coated and uncoated silicone pre-vascularized groups were monitored for up to 90 days (**Fig. 5B**). Animals in both groups remained consistently normoglycemic for up to 70 days, at which time grafts began to destabilize with fluctuating non-fasting blood glucose levels around 11.1 mM (**Fig. 5B – blue line**).

Surprisingly, animals from each of the three groups responded to a glucose bolus when challenged with an OGTT at day 14, returning to normoglycemia within 120 minutes post-administration (**Fig. 5D**). There were no significant differences among the three groups in response to the glucose bolus (**Fig. 5E**).

Similar to the SCID/bg mice, retrieving the grafts for histological analysis was more challenging than anticipated. Islet grafts from the unmodified subcutaneous group did not have any insulin positive staining, consistent with the poor functional outcome observed on normalizing blood glucose levels. At day 21, we were able to retrieve 4 of 8 grafts from animals within the MAA-coated pre-vascularization group. Grafts that were retrieved at day 21 showed extensive insulin staining with intact pancreatic islets and intra-islet vasculature (**Fig. 6B**). Both intra-islet vessels and the vessels throughout the surrounding area were mature (CD31^+^ and SMA^+^). The average CD31^+^ vessel density of the explanted grafts (islet and surrounding tissue) was 178 ± 8 vessels per mm^2^ (**Fig. 6C**). Grafts at day 90 were unable to be reliably retrieved since they were completely integrated with the host’s subcutaneous space.

**Figure 6:**
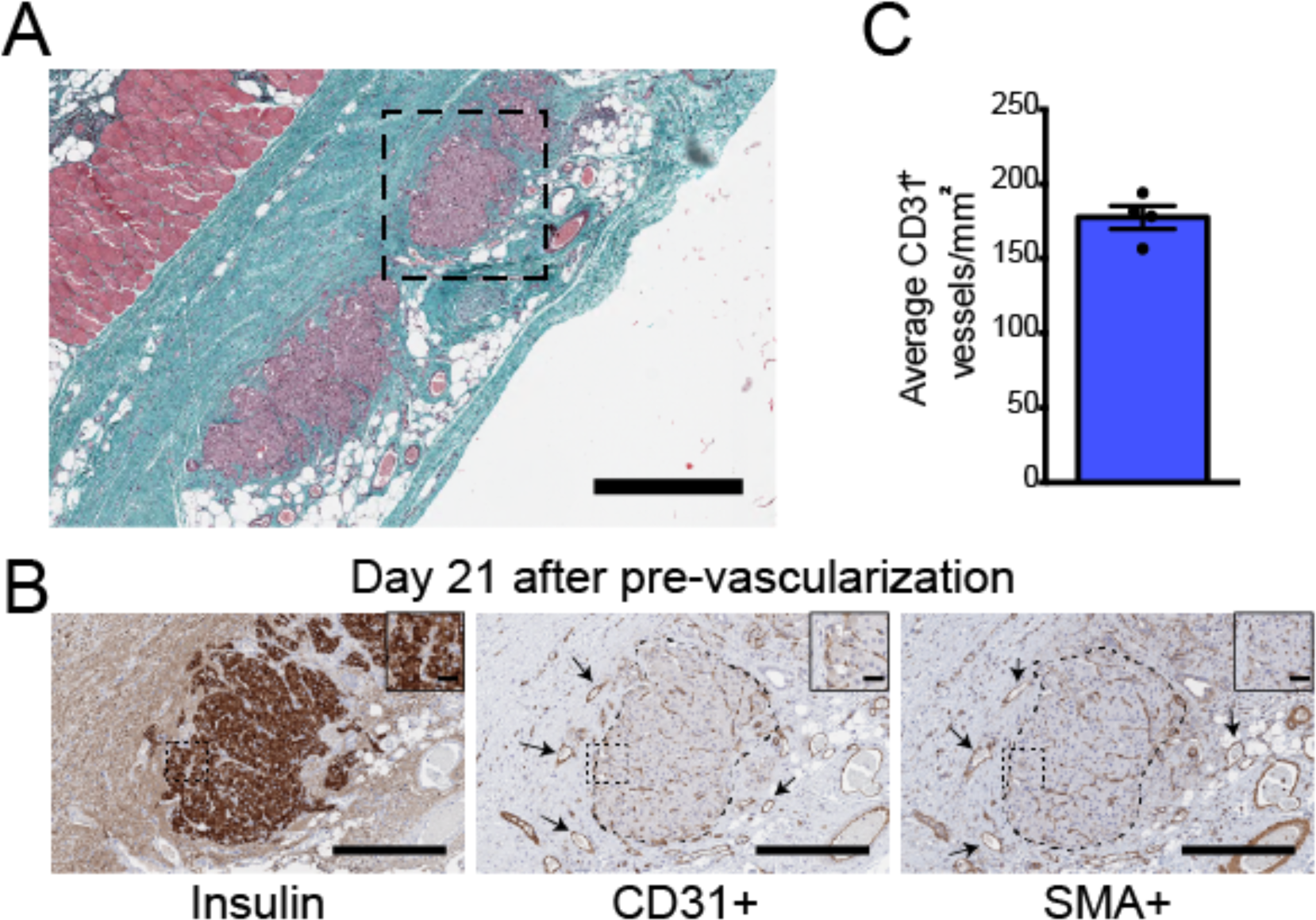
Pancreatic islets recovered from the pre-vascularized space created with the MAA-coated tube in SD rats. **A)** Representative histological image of pancreatic islets at day 21 transplanted into the pre-vascularized space created by the MAA-coated tube. 700 μm scale bar. The pancreatic islets were **B)** insulin^+^ with both mature intra-islet and peripheral vessels (CD31^+^ and SMA^+^). Scale bars of main images = 200 μm and the scale bars of the insets = 50 μm. **C)** While only four of the eight grafts could be retrieved, the average number of CD31^+^ vessels of the pancreatic islet grafts (islets + surrounding tissue) in these 4 at day 21 was 180 +/- 8. Average +/- SEM.

## 4.0 Discussion

### 4.1 Using therapeutic polymers to drive vascularization of the subcutaneous space

MAA-based biomaterials have previously been used in several mouse models to generate mature vasculature in the subcutaneous space^3,10–12^, without the need of exogenous growth factors, or cells. In comparison, cell-based approaches for subcutaneous islet transplantation may be affected by inconsistencies in fabrication and vascularization that are highly dependent on the cells used for the given model^1,6,15^. Furthermore, the co-delivery of endothelial cells^1,4^ (or other vascularizing cells) may require the use of a more potent immunosuppression protocol, since many of these cells actively interact with the host’s immune system^16^.

To reduce the therapeutic dose required to achieve normoglycemia in the subcutaneous space, pre-vascularization is a suitable alternative where vessels are generated before islet transplantation to provide the necessary oxygen and nutrients to support islet engraftment and survival^5,7,9^. We hypothesized that a purely biomaterial-based approach for pre-vascularization would be simpler to translate than an approach based on using cells to generate vessels^1^. Most importantly, since MAA-based pre-vascularization is not dependent on cellular components, immunosuppression is not required during pre-vascularization. It should be noted that immunosuppression was still required when transplanting allogeneic islets (**Supplementary Fig. 1**), but reducing the amount of systemic immunosuppression is beneficial since prolonged immunosuppression can have toxic secondary effects on the recipient^17^.

Previous studies by Pepper et al^2,8,18^ used nylon or silicone catheters to harness the foreign body response^19,20^ to generate blood vessels in the subcutaneous space for islet transplantation. This type of vascularization was affected by the strength of the host’s innate immune response, and it was observed that C57BL6/J mice, with a more aggressive innate immunity response^21^, resulted in a faster return to normoglycemia compared to Balb/C mice^2,8^. Consistent with these results, we observed that uncoated silicone tubes generated many vessels in the subcutaneous space of both mice and rats, presumably by generating a foreign body response that was halted by removing the tube at day 14 (**Fig. 2B, 4B**).

We followed a protocol similar to that used by Pepper et al but coated the silicone catheter with polyMAA-co-IDA. We have previously shown that MAA based materials activate the insulin growth factor (IGF-1)^11^ and sonic hedgehog (Shh)^10^ signalling pathways to promote macrophage polarization towards a M2-like phenotype and drive angiogenesis^12^. This active angiogenic effects from the MAA-coating accounts for the significant increase in the number of vessels compared to the uncoated silicone tube controls in both mice and rats (**Fig. 2B, 4A**). It should be noted that vessels generated with MAA (in the absence of transplanted cells) eventually regress, and by day 21 the number of vessels declined compared to day 14, which was consistent with tissue remodeling as shown previously^11,12^.

### 4.2 Pre-vascularization and islet transplantation

In the streptozotocin-induced diabetic SCID/bg model, a marginal dose of pancreatic islets (250 IEQ) showed a greater therapeutic efficacy when transplanted into the pre-vascularized cavity (both MAA-coated and uncoated silicone groups) compared to transplantation into the unmodified space alone (**Fig. 3A**). In previous studies without pre-vascularization (but with a similar number of endothelial cells), the therapeutic dosage required to achieve normoglycemia was approximately 750 IEQ^1,3^. Although one animal in the unmodified subcutaneous space group (no pre-vascularization) returned to normoglycemia, there was a significant delay in the time it took to achieve normoglycemia (**Fig. 3B**). Pancreatic islets that survive the immediate hypoxic conditions secrete VEGF-A^22^ to recruit neutrophils^23^ and drive re-vascularization once transplanted, which may explain the delay observed in the no pre-vascularization group relative to the groups with pre-vascularization (**Fig. 3A, B**).

The benefits of MAA-driven pre-vascularization were most apparent in the larger SD rat model (**Fig. 5A**). MAA-coated tubes generated the greatest number of vessels (**Fig. 4B**) and showed the most consistent return to normoglycemia (**Fig. 5A**), compared to the less vascularized site associated with the uncoated silicone tube group (**Fig. 4B**). Due to the avascular nature of the subcutaneous space, 4000 IEQ did not have any therapeutic effect when transplanted into the unmodified subcutaneous space (**Fig. 5A – red line**). Interestingly, all groups could respond to an OGTT at day 14 (**Fig. 5D**), suggesting that although islet grafts could not regulate non-fasting blood glucose levels, some residual islet function remained sufficient to respond in a glucose challenge test^24^.

The co-delivery of HUVEC with the islets in SCID/bg (but not the rat) may have also contributed to generating a more robust vascularization response after transplantation into the pre-vascularized site. In previous studies, the addition of HUVEC with islet collagen microtissues generated a greater number of vessels at early time points and prevented collagen degradation compared to grafts without HUVEC following subcutaneous islet transplantation^1^. HUVEC were included to enable a comparison with these previous studies where 750 IEQ was needed to restore normoglycemia without pre-vascularization. HUVEC help drive re-vascularization of transplanted pancreatic islets by secreting VEGF^25^ (in addition to that produced by islets) and other pro-angiogenic growth factors^26^. However, by co-delivering HUVEC, the perceived therapeutic benefit of having more vessels before islet transplantation (**Fig. 2A, B**) may have been obscured.

Retrieval of the islet grafts was more challenging than anticipated in the SD rat model, unlike what was seen in the SCID/bg mouse with or without^1^ pre-vascularization. In the rat, pancreatic islets were delivered within a collagen gel to support their engraftment and function^27–29^, however collagen degrades over time when subcutaneously transplanted without endothelial cells^1^. In our explants, we could only find intact islets (insulin^+^) in implants that were harvested at day 21 in the MAA-coated tube group (**Fig. 6B)**. The area surrounding the embedded islets was well vascularized (CD31^+^ and SMA^+^), and the presence of intra-islet vessels was also observed (**Fig. 6B, C**). Explanted grafts from the uncoated tube or unmodified subcutaneous groups could not be distinguished from the surrounding subcutaneous space. There was little or no insulin staining in the explanted tissue and may be due to poor islet survival, since the uncoated tube and unmodified subcutaneous space groups had significantly higher non-fasting blood glucose levels compared to the MAA-coated tube group (**Fig. 5A**).

At day 90, grafts had completely integrated with the host’s tissue and could not be distinguished from the surrounding subcutaneous space making retrievability not possible. Although we did not directly confirm islet integration and re-vascularization using CLARITY as in previous studies^1,4^, we expect that the vessel islet integration seen before^1^ was critical for their therapeutic efficacy. Long-term integration of the islet graft was possible because the collagen delivery vehicle was remodeled, but this integration can make retrieval more difficult, unlike approaches delivering islets within a device^5^. Future experiments could use image-guided surgery^30^ to reliably remove the graft (without sacrificing the animal) after incorporating fluorescently labeled gold or other nanoparticles into the collagen gel^31^.

### 4.3 Considerations for translating pre-vascularization approaches to larger, immune-competent models

To model the clinical allograft transplantation scenario, it was necessary to use systemic immunosuppression to overcome the adaptive immune response^32^. Unlike previous studies transplanting allogeneic islets in the omentum of SD rats^33^, tacrolimus alone did not prevent rapid graft failure in the pre-vascularized subcutaneous space (**Supplementary Figure 3A** – **green line**). It has been previously reported that there are site-specific differences in the levels of immunosuppressive drugs^34^ and kinetics of allo-antigen specific immunity^35^, which can affect islet engraftment. Tacrolimus was sufficient to enable allogeneic rat endothelial cell transplants^36^ in the subcutaneous space but not islet transplants. In our study, we found a combination of ALS, fingolimod and MMF was necessary to prevent rapid graft rejection (**Fig. 5A**).

Tacrolimus is a calcineurin inhibitor, which inhibits T-cell function, but its immunosuppressive effect can be variable and is dependent on its bioavailability^37^, which may be dependent on the transplant site. In comparison, a combination of anti-lymphocyte serum, fingolimod and mycophenolic acid provided consistent immunosuppression in others’ studies^38^. Anti-lymphocyte serum contains antibodies that block the activation of host lymphocytes, fingolimod prevents the sequestration of lymphocytes from the lymph nodes^39^ and mycophenolic acid inhibits T-cell and B-cell function^40^. This combination protocol is preferred because increasing the dosage of tacrolimus can be detrimental to islet health and function^41^.

Scaling up these approaches into humans is challenging due to the large transplant volume required to deliver a sufficient number of islets (approximately 800,000 IEQ) to achieve insulin independence (∼10,000 – 12,000 IEQ/kg)^42,43^. Assuming a 5-10% packing volume, the mean packing volume used to deliver this number of pancreatic islets is approximately 15-30 mL. Other alternative sites such as the kidney capsule (normally used in pre-clinical research^44,45^) is not clinically feasible due to the limited available volume of the site^46^. In the approach used here, the volume of the pre-vascularized space was scalable by changing the length of tubing. For example, by using an MAA coated tube (3.18 mm OD) with a length of 10 cm the theoretical available volume for transplantation is ∼0.8 mL, enough for ∼30,000 IEQ at a density of 37,500 IEQ/mL (6.4% v/v loading as in the rat studies). To scale this approach for human transplants and to achieve for example, a volume of 15-30 mL multiple longer tubes can be implanted in parallel to increase the available transplant volume; this corresponds to approximately 100 cm^2^ in area, depending on spacing between parallel tubes.

Using MAA-coated tubes, the pre-vascularized subcutaneous space is a feasible alternative site for subcutaneous islet transplantation, which we believe can be used to support a scaled-up transplant volume in larger animal models and humans, while minimizing the invasiveness associated with portal vein infusion. This remains to be tested; however, we anticipate the vascularizing activity of MAA to be present in animals other than rodents, since the relevant pathways (e.g. Shh and IGF-1) are highly conserved among different species^47,48^.

Some grafts eventually failed at the >70 day point as the recipients outgrew their implant (**Fig. 5B**). At the time of transplantation, SD rats used as recipients had an average weight of 319 ± 49 g (mean ± SD, **Supplementary Figure 4A**) and were transplanted with 4000 IEQ, resulting in a dosage of approximately 13 × 10^3^ ± 2 × 10^3^ IEQ/kg (mean ± SD, **Supplementary Figure 4B**). At the time of transplantation this dose was comparable to the clinically relevant dose used in the portal vein^42^. However, the recipients continued to gain weight and by day 70 their average weight was 561 ± 9; thus the islet dose was nominally only 7 × 10^3^ IEQ by the end of the experiment – **Supplementary Fig. 4C**). Future long-term studies using fully grown (i.e., older) rats or pigs could be used to reduce any confounding variable that weight gain may have on graft function over time.

## 5.0 Conclusion

In this study we used a MAA-based coating of a silicone tube to form a pre-vascularized cavity for the transplantation of pancreatic islets. The MAA coating drove a more robust vascularization response compared to the uncoated silicone tube control and this was important for islet transplantation efficacy in larger rodent models. A marginal mass of pancreatic islets transplanted into the pre-vascularized space with MAA was sufficient to return streptozotocin-induced diabetic SCID/bg mice to normal. More importantly this approach was scalable to a larger rat transplant model and a clinically relevant therapeutic dosage was sufficient in returning diabetic rats to normal. Using a suitable immunosuppression protocol, allogeneic transplanted pancreatic islets survived in the pre-vascularized subcutaneous space and became revascularized. Islets transplanted in the unmodified subcutaneous space did not have any effect on returning animals to normoglycemia. This work highlights an approach that has the potential to be scaled into larger animal models, such as pigs or non-human primates without the need for additional cells, or growth factors to pre-vascularize the subcutaneous space to support islet transplantation.

## Acknowledgements

We would like to thank Chuen Lo for his help and expertise with animal surgeries and animal isolation. We would also like to thank Dr. Cherie Stabler (University of Florida) for kindly sharing the immunosuppression regimen used in this work. This work was supported by the CIHR (#341676) and the JDRF (3-SRA-2016-253-S-B). A.E.V. was supported by scholarships from the Province of Ontario, the Jennifer Dorrington Award, and the Banting and Best Diabetes Centre.

## Author Contributions

A.E.V. and M.V.S. conceived the study. A.E.V., I.T., S.M.K. performed the experiments for the manuscript. A.E.V. and I.T. analyzed the data for the manuscript. A.E.V., and M.V.S. wrote the manuscript. All of the authors edited and provided feedback for the manuscript.

## Declaration of Interests

The authors declare no competing interests.

## Supplementary Figures

**Supplementary Figure 1:**
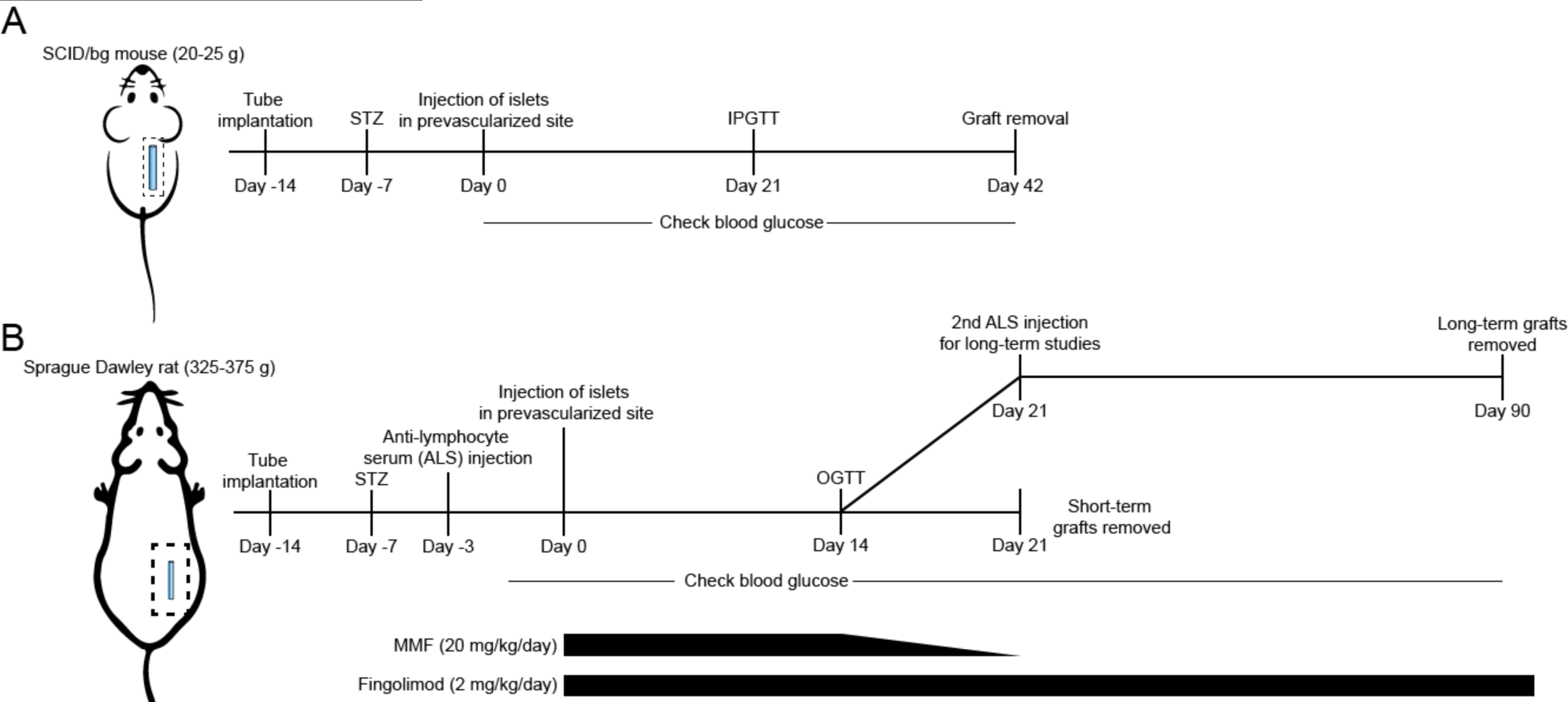
Experimental setup of pre-vascularization and islet transplantation experiments. **A)** SCID/bg mice were subcutaneously implanted with either an MAA-coated silicone tube or an uncoated silicone tube or left unmodified. 7 days prior to transplantation, the mice were made diabetic using a single i.p. injection of streptozotocin. At time of transplantation (day 0), the tubes were removed, and islets embedded within a collagen gel with HUVEC were implanted into the pre-vascularized site or an unmodified subcutaneous space. Non-fasting blood glucose readings were checked daily and IPGTT was administered on day 21. Normoglycemic animals were sacrificed at day 42 and their explants were assessed via histology; unmodified animals were sacrificed at day 28. **B)** The implantation of the MAA-coated tube into the subcutaneous space of Sprague Dawley rats was identical to the SCID/bg mouse. However, on day 3 before islet transplantation, the rats were administered a dose of anti-lymphocyte serum. On the day of the allogeneic islet transplantation (after the initial 14 days), animals were given a daily oral gavage with mycophenolic acid (20 mg/kg/day, that was tapered down to 0 mg/kg between days 14 to 21) and fingolimod (2 mg/kg/day). Non-fasting blood glucose readings were measured daily and an OGTT was administered on day 14 after islet transplantation. On day 21, animals for the short-term studies were sacrificed, and their grafts were assessed via histology. For long-term studies, an additional injection of ALS was given at day 21 and animals were tracked until day 90, when their grafts were removed.

**Supplementary Figure 2:**
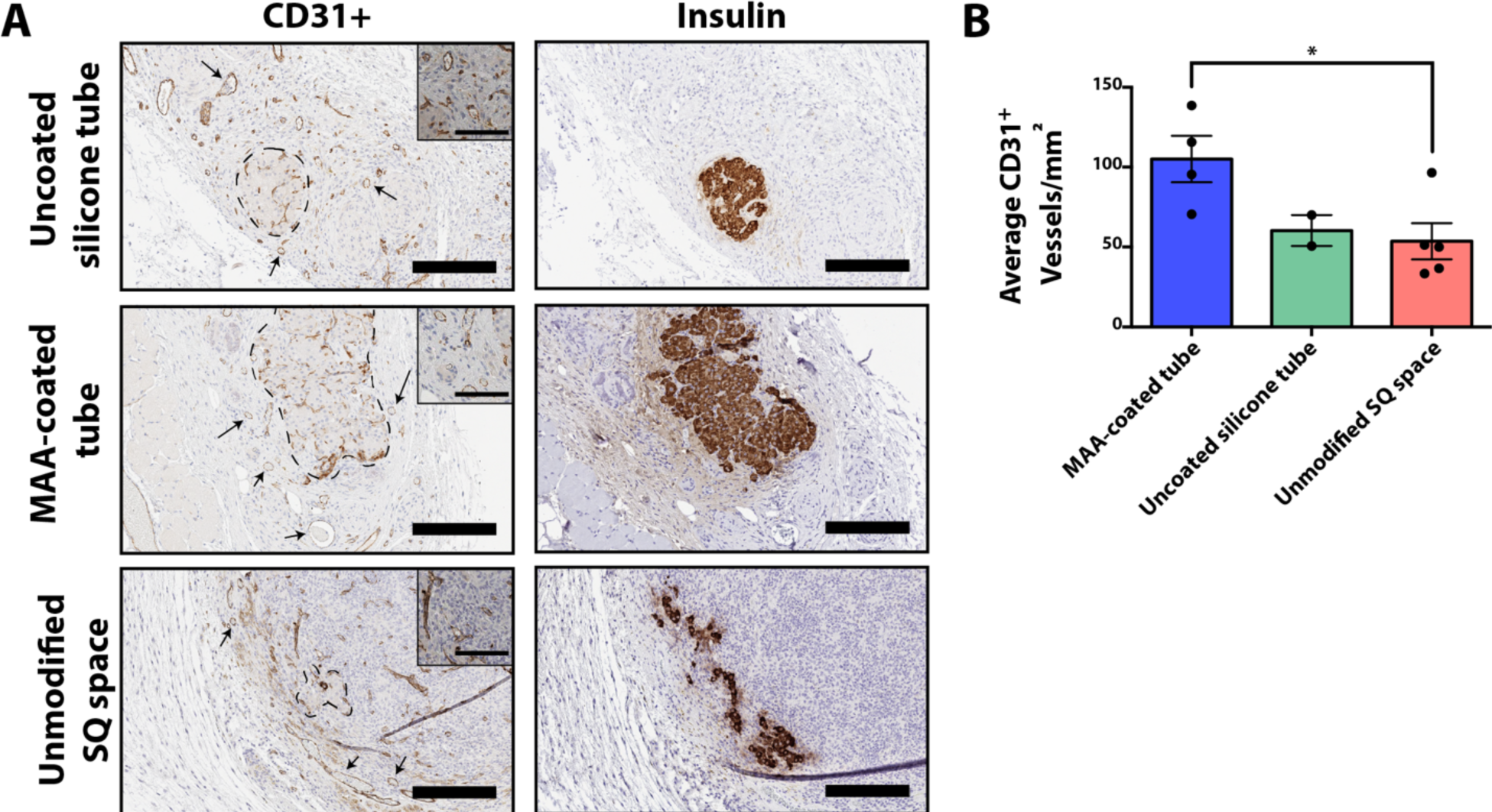
Pancreatic islets recovered from the pre-vascularized space of SCID/bg mice. **A)** Pancreatic islets delivered into the pre-vascularized subcutaneous space were vascularized with intra-islet vessels (CD31^+^ within dotted line) and were intact (insulin^+^) at day 42. However, delivery into the unmodified subcutaneous space resulted in fragmented islets that did not have intra-islet vessels clearly present (CD31^+^) at day 28. CD31^+^ vessels were present throughout the collagen + EC grafts in all three groups. Arrows denote examples of CD31^+^ vessels. Scale bars = 200 μm. **B)** There was a significantly greater CD31^+^ vessel density in the grafts transplanted into the pre-vascularized space using the MAA-coated tube compared to the unmodified subcutaneous space (p < 0.05). However, only 4 of 5 grafts in the MAA-coated group, 2 of 5 grafts in the uncoated silicone tube group could be reliably explanted from the subcutaneous space. Average +/- SEM, n = 4 for MAA-coated group, n = 2 for uncoated group, and n = 5 for unmodified SQ space group, *p < 0.05.

**Supplementary Figure 3:**
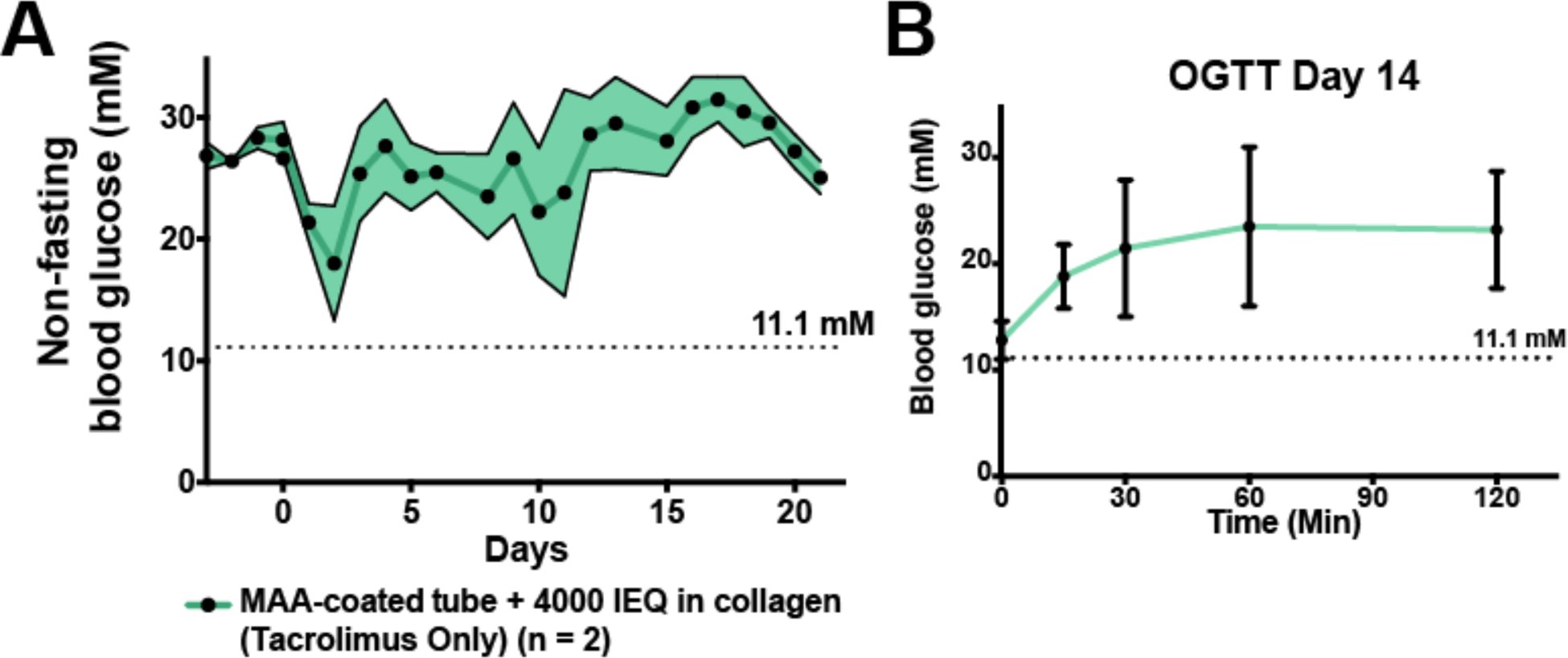
Tacrolimus alone was insufficient immunosuppression for allogeneic islet transplantation in the MAA coated tube pre-vascularized group. **A)** Average non-fasting blood glucose levels of animals transplanted with 4000 IEQ into the pre-vascularized subcutaneous space created with a MAA-coated tube. n = 2 **B)** Animals fasting blood glucose levels were 11.1 mM after overnight fasting, but animals were intolerant to an oral glucose tolerance test on day 14.

**Supplementary Figure 4:**
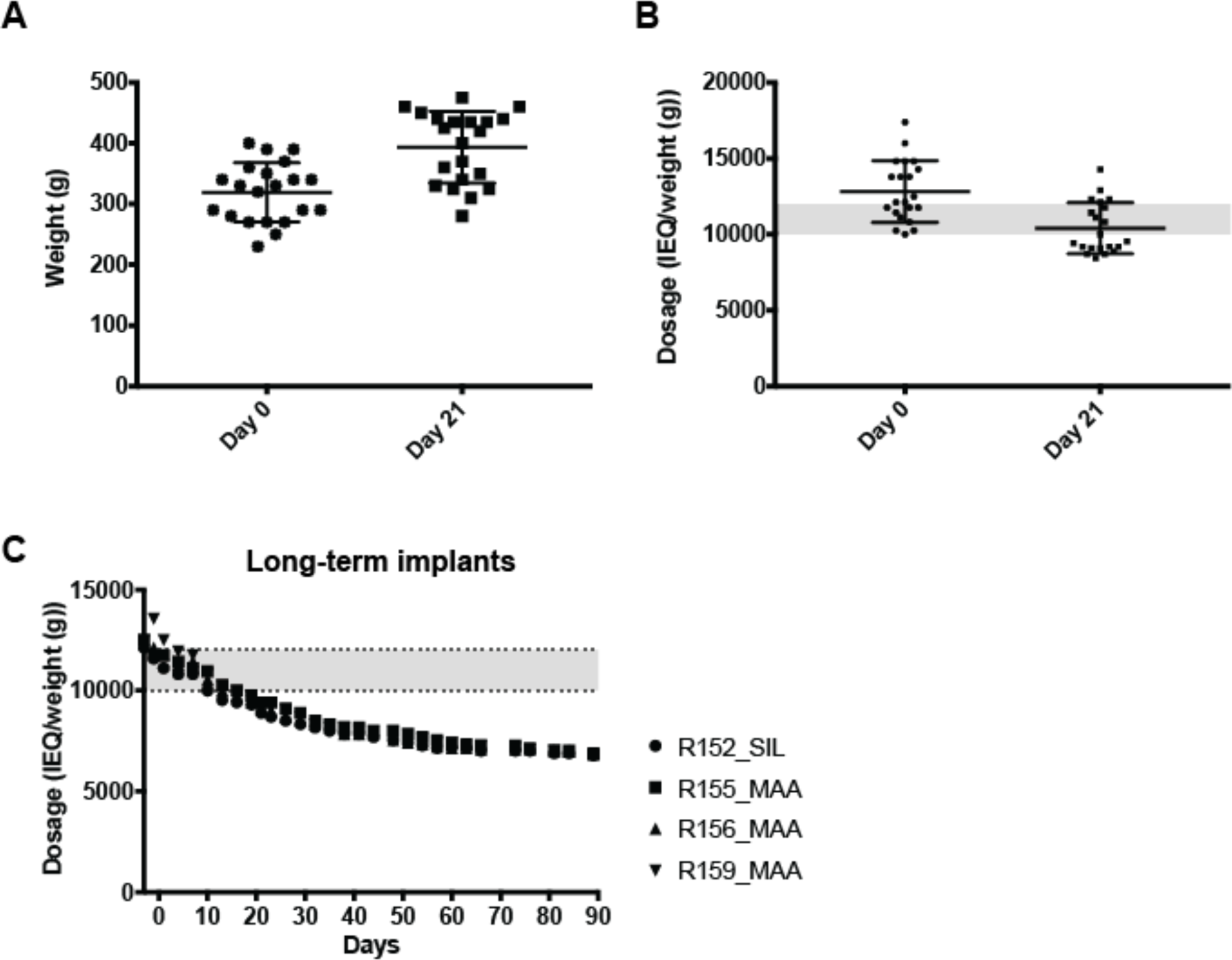
Changes in weight over time for SD rats subcutaneously transplanted with allogeneic islets. **A)** The average weight of recipients used in the allogeneic SD rat study at days 0 and 21. **B)** SD rats were transplanted with 4000 allogeneic IEQ at day 0. The apparent transplant dose was calculated by taking the number of transplanted IEQ (4000) and dividing it by the weight of each recipient. The shaded grey region represents the effective dosage transplanted into the portal vein of humans to achieve insulin independence (10,000 – 12,000 IEQ/kg)^42^ **C)** The apparent dose received by recipients with long-term implants over time. Animals had apparent doses lower than the clinically effective levels starting at day 21 and continued to drop as animals gained weight.

**Supplementary Figure 5:**
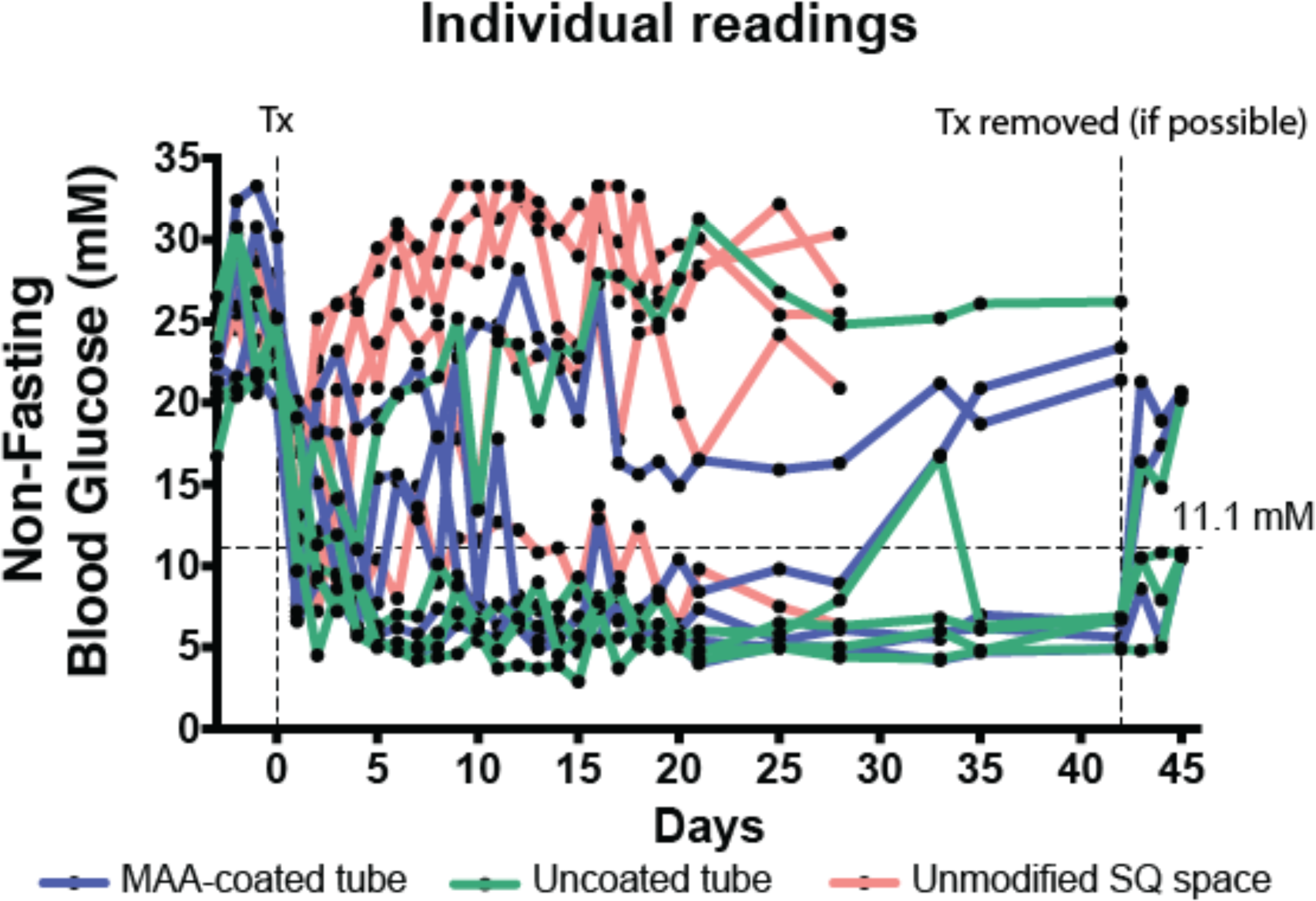
Individual non-fasting blood glucose readings of SCID/bg mice transplanted with 250 IEQ mixed with collagen and HUVEC. After 28 days, grafts were removed from the unmodified subcutaneous group. After 42 days, grafts in the MAA, and uncoated groups were explanted without sacrificing the animals, however some explants (1 of 3 for MAA, and 3 of 4 for uncoated) could not be fully removed.

## Supplementary Tables

**Supplementary Table 1:** The theoretical available transplant volume is dependent on the length of tubing used to pre-vascularize the subcutaneous space. For humans, multiple tubes can be implanted in parallel to increase the effective transplant volume.

| Tube diameter (cm) | Tube Length (cm) | Theoretical Available volume (mL) |
| --- | --- | --- |
| 0.318 cm | 3 cm | 0.24 mL |
| 0.318 cm | 6 cm | 0.48 mL |
| 0.318 cm | 10 cm | 0.79 mL |

## Notes

### Competing Interest Statement

The authors have declared no competing interest.

## References

1. Vlahos, A. E., Cober, N. & Sefton, M. V. Modular tissue engineering for the vascularization of subcutaneously transplanted pancreatic islets. Proc. Natl. Acad. Sci. USA 114, 9337–9342 (2017).

2. Pepper, A. R. et al. A prevascularized subcutaneous device-less site for islet and cellular transplantation. Nat. Biotechnol. 33, 518–523 (2015).

3. Mahou, R., Zhang, D. K. Y., Vlahos, A. E. & Sefton, M. V. Injectable and inherently vascularizing semi-interpenetrating polymer network for delivering cells to the subcutaneous space. Biomaterials 131, 27–35 (2017).

4. Vlahos, A. E. et al. Endothelialized collagen based pseudo-islets enables tuneable subcutaneous diabetes therapy. Biomaterials 232, 119710 (2020).

5. Pepper, A. R. et al. Diabetes is reversed in a murine model by marginal mass syngeneic islet transplantation using a subcutaneous cell pouch device. Transplantation 99, 2294–2300 (2015).

6. Song, W. et al. Engineering transferrable microvascular meshes for subcutaneous islet transplantation. Nat. Commun. 10, 4602 (2019).

7. Smink, A. M. et al. The Efficacy of a Prevascularized, Retrievable Poly(D,L,-lactide-co-ε-caprolactone) Subcutaneous Scaffold as Transplantation Site for Pancreatic Islets. Transplantation 101, e112–e119 (2017).

8. Pepper, A. R. et al. Harnessing the Foreign Body Reaction in Marginal Mass Device-less Subcutaneous Islet Transplantation in Mice. Transplantation 100, 1474–1479 (2016).

9. Luan, N. M. & Iwata, H. Long-term allogeneic islet graft survival in prevascularized subcutaneous sites without immunosuppressive treatment. Am. J. Transplant. 14, 1533–1542 (2014).

10. Lisovsky, A., Zhang, D. K. Y. & Sefton, M. V. Effect of methacrylic acid beads on the sonic hedgehog signaling pathway and macrophage polarization in a subcutaneous injection mouse model. Biomaterials 98, 203–214 (2016).

11. Talior-Volodarsky, I., Mahou, R., Zhang, D. & Sefton, M. The role of insulin growth factor-1 on the vascular regenerative effect of MAA coated disks and macrophage-endothelial cell crosstalk. Biomaterials 144, 199–210 (2017).

12. Coindre, V. F., Carleton, M. M. & Sefton, M. V. Methacrylic acid copolymer coating enhances constructive remodeling of polypropylene mesh by increasing the vascular response. Adv Healthc Mater 8, e1900667 (2019).

13. Coindre, V., Kinney, S. M. & Sefton, M. V. Methacrylic Acid Copolymer Coating of Polypropylene Mesh Chamber Improves Subcutaneous Islet Engraftment. (2020).

14. Wells, L. A. & Sefton, M. V. The effect of methacrylic acid in smooth coatings on dTHP1 and HUVEC gene expression. Biomater. Sci. 2, 1768–1778 (2014).

15. Kim, J.-S. et al. Vascularization of PLGA-based bio-artificial beds by hypoxia-preconditioned mesenchymal stem cells for subcutaneous xenogeneic islet transplantation. Xenotransplantation 26, e12441 (2019).

16. Mai, J., Virtue, A., Shen, J., Wang, H. & Yang, X.-F. An evolving new paradigm: endothelial cells--conditional innate immune cells. J Hematol Oncol 6, 61 (2013).

17. Shaw, L. M., Kaplan, B. & Kaufman, D. Toxic effects of immunosuppressive drugs: mechanisms and strategies for controlling them. Clin. Chem. 42, 1316–1321 (1996).

18. Pepper, A. R. et al. Transplantation of human pancreatic endoderm cells reverses diabetes post transplantation in a prevascularized subcutaneous site. Stem Cell Rep. 8, 1689–1700 (2017).

19. Anderson, J. M., Rodriguez, A. & Chang, D. T. Foreign body reaction to biomaterials. Semin. Immunol. 20, 86–100 (2008).

20. Jones, J. A. et al. Proteomic analysis and quantification of cytokines and chemokines from biomaterial surface-adherent macrophages and foreign body giant cells. J. Biomed. Mater. Res. A 83, 585–596 (2007).

21. King, A., Sandler, S. & Andersson, A. The effect of host factors and capsule composition on the cellular overgrowth on implanted alginate capsules. J. Biomed. Mater. Res. 57, 374–383 (2001).

22. Brissova, M. et al. Pancreatic islet production of vascular endothelial growth factor--a is essential for islet vascularization, revascularization, and function. Diabetes 55, 2974–2985 (2006).

23. Christoffersson, G. et al. VEGF-A recruits a proangiogenic MMP-9-delivering neutrophil subset that induces angiogenesis in transplanted hypoxic tissue. Blood 120, 4653–4662 (2012).

24. Dy, E. C., Harlan, D. M. & Rother, K. I. Assessment of islet function following islet and pancreas transplantation. Curr Diab Rep 6, 316–322 (2006).

25. Gerhardt, H. et al. VEGF guides angiogenic sprouting utilizing endothelial tip cell filopodia. J. Cell Biol. 161, 1163–1177 (2003).

26. Ucuzian, A. A., Gassman, A. A., East, A. T. & Greisler, H. P. Molecular mediators of angiogenesis. J Burn Care Res 31, 158–175 (2010).

27. Stephens, C. H. et al. In situ type I oligomeric collagen macroencapsulation promotes islet longevity and function in vitro and in vivo. Am. J. Physiol. Endocrinol. Metab. 315, E650–E661 (2018).

28. Hammar, E. et al. Extracellular matrix protects pancreatic beta-cells against apoptosis: role of short- and long-term signaling pathways. Diabetes 53, 2034–2041 (2004).

29. Zhao, Y. et al. Preservation of islet survival by upregulating α3 integrin signaling: the importance of 3-dimensional islet culture in basement membrane extract. Transplant. Proc. 42, 4638–4642 (2010).

30. Nguyen, Q. T. & Tsien, R. Y. Fluorescence-guided surgery with live molecular navigation--a new cutting edge. Nat. Rev. Cancer 13, 653–662 (2013).

31. Castaneda, L., Valle, J., Yang, N., Pluskat, S. & Slowinska, K. Collagen cross-linking with Au nanoparticles. Biomacromolecules 9, 3383–3388 (2008).

32. Brehm, M. A. et al. Human immune system development and rejection of human islet allografts in spontaneously diabetic NOD-Rag1null IL2rgammanull Ins2Akita mice. Diabetes 59, 2265–2270 (2010).

33. Gupta, R. & Sefton, M. V. Application of an endothelialized modular construct for islet transplantation in syngeneic and allogeneic immunosuppressed rat models. Tissue Eng. Part A 17, 2005–2015 (2011).

34. Cantarelli, E. et al. Murine animal models for preclinical islet transplantation: No model fits all (research purposes). Islets 5, 79–86 (2013).

35. Wood, K. J. & Goto, R. Mechanisms of rejection: current perspectives. Transplantation 93, 1–10 (2012).

36. Chamberlain, M. D., Gupta, R. & Sefton, M. V. Chimeric vessel tissue engineering driven by endothelialized modules in immunosuppressed Sprague-Dawley rats. Tissue Eng. Part A 17, 151–160 (2011).

37. Kapturczak, M. H., Meier-Kriesche, H. U. & Kaplan, B. Pharmacology of calcineurin antagonists. Transplant. Proc. 36, 25S–32S (2004).

38. Buchwald, P. et al. Feasibility of localized immunosuppression: 1. Exploratory studies with glucocorticoids in a biohybrid device designed for cell transplantation. Pharmazie 65, 421–428 (2010).

39. Budde, K. et al. FTY720 (fingolimod) in renal transplantation. Clin Transplant 20 Suppl 17, 17–24 (2006).

40. Shaw, L. M., Mick, R., Nowak, I., Korecka, M. & Brayman, K. L. Pharmacokinetics of mycophenolic acid in renal transplant patients with delayed graft function. J Clin Pharmacol 38, 268–275 (1998).

41. Paty, B. W., Harmon, J. S., Marsh, C. L. & Robertson, R. P. Inhibitory effects of immunosuppressive drugs on insulin secretion from HIT-T15 cells and Wistar rat islets. Transplantation 73, 353–357 (2002).

42. Shapiro, A. M. et al. Islet transplantation in seven patients with type 1 diabetes mellitus using a glucocorticoid-free immunosuppressive regimen. N. Engl. J. Med. 343, 230–238 (2000).

43. Ryan, E. A. et al. Five-year follow-up after clinical islet transplantation. Diabetes 54, 2060–2069 (2005).

44. Morini, S. et al. Revascularization and remodelling of pancreatic islets grafted under the kidney capsule. J. Anat. 210, 565–577 (2007).

45. Szot, G. L., Koudria, P. & Bluestone, J. A. Transplantation of pancreatic islets into the kidney capsule of diabetic mice. J. Vis. Exp. 404 (2007). doi: 10.3791/404

46. Cantarelli, E. & Piemonti, L. Alternative transplantation sites for pancreatic islet grafts. Curr Diab Rep 11, 364–374 (2011).

47. Barbieri, M., Bonafè, M., Franceschi, C. & Paolisso, G. Insulin/IGF-I-signaling pathway: an evolutionarily conserved mechanism of longevity from yeast to humans. Am. J. Physiol. Endocrinol. Metab. 285, E1064–71 (2003).

48. Wilson, C. W. & Chuang, P.-T. Mechanism and evolution of cytosolic Hedgehog signal transduction. Development 137, 2079–2094 (2010).

